# Machine learning-predicted chromatin organization landscape across pediatric tumors

**DOI:** 10.1101/2025.03.28.645984

**Authors:** Ketrin Gjoni, Shu Zhang, Rachel E. Yan, Bo Zhang, Daniel Miller, Adam Resnick, Nadia Dahmane, Katherine S. Pollard

**Author notes:** denotes equal contribution (Ketrin Gjoni, Shu Zhang).

## Abstract

Structural variants (SVs) are increasingly recognized as important contributors to oncogenesis through their effects on 3D genome folding. Recent advances in whole-genome sequencing have enabled large-scale profiling of SVs across diverse tumors, yet experimental characterization of their individual impact on genome folding remains infeasible. Here, we leveraged a convolutional neural network, Akita, to predict disruptions in genome folding caused by somatic SVs identified in 61 tumor types from the Children’s Brain Tumor Network dataset. Our analysis reveals significant variability in SV-induced disruptions across tumor types, with the most disruptive SVs coming from lymphomas and sarcomas, metastatic tumors, and germline cell tumors. Dimensionality reduction of disruption scores identified five recurrently disrupted regions enriched for high-impact SVs across multiple tumors. Some of these regions are highly disrupted despite not being highly mutated, and harbor tumor-associated genes and transcriptional regulators. To further interpret the functional relevance of high-scoring SVs, we integrated epigenetic data and developed a modified Activity-by-Contact scoring approach to prioritize SVs with disrupted genome contacts at active enhancers. This method highlighted highly disruptive SVs near key oncogenes, as well as novel candidate loci potentially implicated in tumorigenesis. These findings highlight the utility of machine learning for identifying novel SVs, loci, and genetic mechanisms contributing to pediatric cancers. This framework provides a foundation for future studies linking SV-driven regulatory changes to cancer pathogenesis.

## Introduction

Pediatric brain tumors (PBTs) are the most common solid cancers in children and have seen little improvement in treatment or understanding compared to other childhood cancers^1^. One reason is that the genetic causes behind many PBTs remain unclear, and efforts are ongoing to dissect the underlying molecular mechanisms that promote tumorigenesis in these cancers.

Structural variants (SVs)–large deletions, duplications, inversions, insertions, and chromosomal rearrangements of genomic regions–are major drivers of cancer that play an important role in shaping the transcriptomic landscape of PBTs^2^. SVs can disrupt tumor suppressors or amplify oncogenes by fusing genes, altering regulatory elements, or directly disrupting the genes themselves. In cancer, SVs have been shown to create harmful gene fusions such as *BCR-ABL* in leukemias and *MYB-QKI* rearrangements in pediatric gliomas^3,4^. Overall, PBTs have higher rates of SVs compared to other pediatric brain tumors^5^ and their SV patterns differ from adult tumors^6^. Several SV signatures converge to alter *MYC*, suggesting the importance of the role of SVs in cancer progression^6^.

More recently, SVs have been suggested to cause cancer through alterations to genome organization^7^. In humans, the genome is folded into higher order structures, ranging from small loops of only a few kilobases (kb) in length to larger topologically associating domains (TADs) that span multiple megabases (Mb) of DNA^8^. By measuring genome organization through conformation capture experiments, the effect of SVs on structures like TADs has been implicated in various PBTs. The deletion of a TAD boundary can merge two TADs and cause genes from one TAD to hijack enhancers from the previously distinct other TAD and become overexpressed. Enhancer hijacking has been shown to lead to the activation of oncogenes across cancer types^9,10^, including pediatric high-grade gliomas^11^ and medulloblastomas^12^. In ependymomas, the SV-caused formation of neo-TADs has been shown to affect essential genes^13^. SVs that result in rearrangements of genomic contacts have been shown to upregulate *TERT* and contribute to neuroblastoma^14,15^. These studies clearly highlight the contribution of SVs to cancer progression through altered genome structure.

We hypothesize that SV-mediated disrupted genome organization contributes to PBTs at large, beyond these individual anecdotal examples. The Children’s Brain Tumor Network (CBTN)–a collaborative and international effort collecting a rich database of pediatric tumor multi-omics and clinical data^16^–allows us to comprehensively study PBT SVs. However, testing the effect of hundreds of thousands of SVs experimentally would be costly, time-consuming, and impossible for larger SVs. To overcome this, we turn to machine learning (ML) to predict 3D genome folding patterns in high throughput and at low cost. The Akita model predicts chromatin contact maps from a ∼1 Mb DNA sequence with high accuracy^17^. This sequence-based model enables the use of *in silico* mutagenesis (ISM) to quantify the isolated effect of individual SVs. SuPreMo-Akita streamlines ISM with Akita by making predictions for DNA sequences with and without an SV of interest and measuring the effect of each SV on the surrounding chromatin contacts^18^. With these tools, we are equipped to systematically test millions of SVs *in silico*, and prioritize ones predicted to be most harmful.

Here, we applied SuPreMo-Akita to nearly 300,000 SVs from pediatric cancer patients in the CBTN cohort to evaluate SV disruption to genome folding and find patterns across and within individual tumor types. We investigated regions that experienced recurrent disruptions to genome organization across tumor types, evaluated variant effects on putative enhancers, and prioritized candidate variants that disrupt tumor-associated genes.

## Results

### Overview of SV profiles across CBTN tumors

We analyzed pediatric tumors from the CBTN cohort for which somatic SVs were called on tumor tissue from 1,843 individuals, at a median of 9.4 years old. The patients were diagnosed with 61 different types of cancer, including combinations of multiple types^16^, which we grouped into 14 broader categories (**Fig. 1, Supplementary Fig. 1, Supplementary Table 1, Methods**). The majority of the samples fell into gliomas (**Fig. 1a**), which include brainstem glioma, low- and high-grade astrocytoma, ganglioglioma, dysembryoplastic neuroepithelial tumor (DNET), gliosis, glial-neuronal tumor, gliomatosis cerebri, subependymal giant cell astrocytoma (SEGA), and oligodendroglioma (**Supplementary Table 1**). Across these, low grade astrocytomas have the highest number of samples in this dataset, with 553 samples (**Supplementary Fig. 1a**). As seen with single nucleotide variant (SNV) mutation rates^19,20^, SV mutation rates also vary across tumor categories. On average, samples had a median of 35 variants, with lymphomas/sarcomas having the highest rates of mutations at a median of 71.5 and mesenchymal tumors, such as Ewing sarcoma and hemangioblastoma, the lowest at 24 variants per sample (**Fig. 1b**).

**Figure 1.**
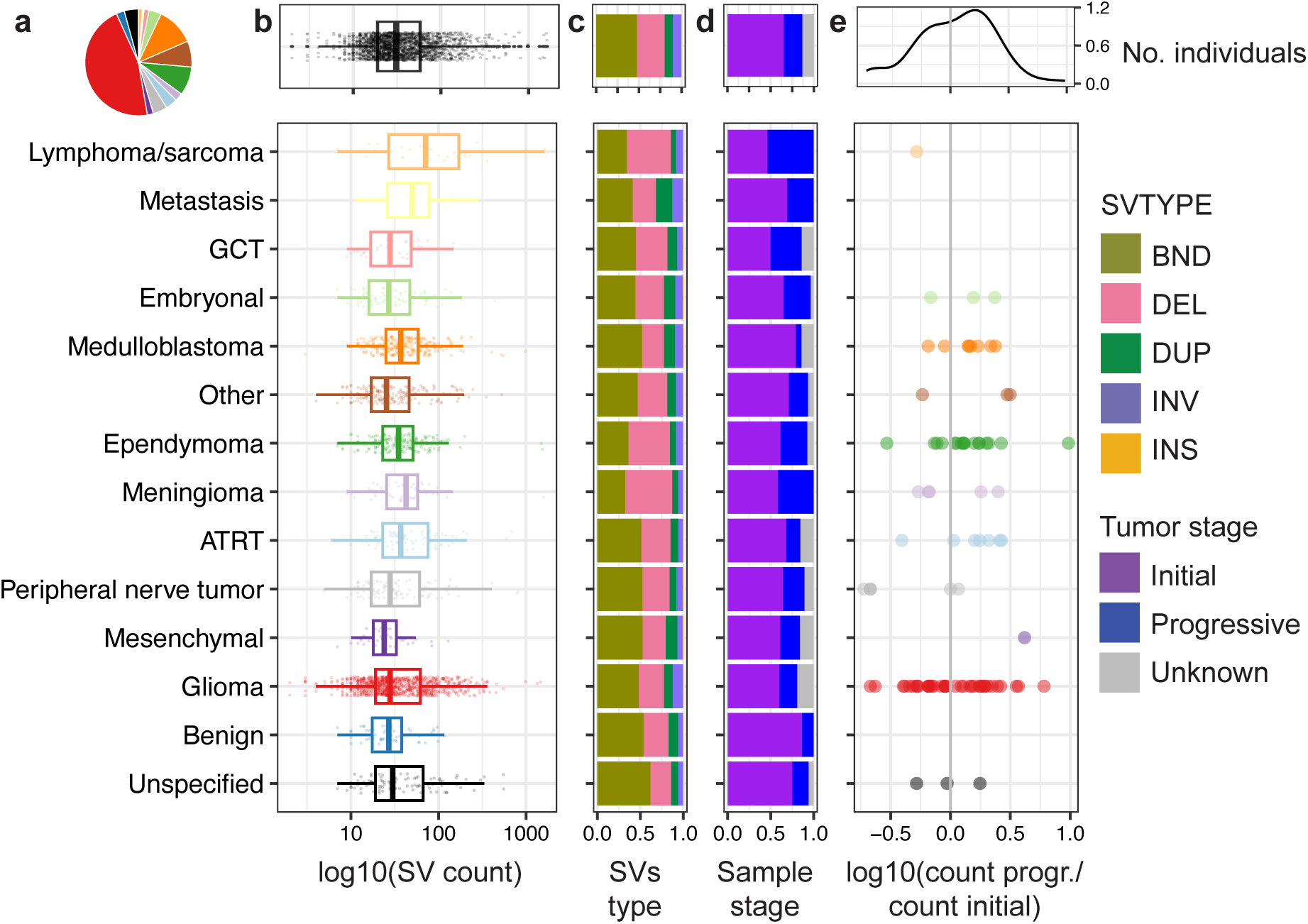
Summary of SV profiles across 14 pediatric tumor categories. **a.** Distribution of the number of tumor samples per tumor category. **b.** Distribution of variant lengths across tumor categories. Each point is a variant, with multiple samples and tumor types included in each category. **c.** Fraction of SV types across tumor categories out of all SVs from all samples in each category. **d.** Fraction of tumor stages per sample out of all samples from all tumor types in each category. **e.** Log fold change in number of variants in progressive over initial samples from each participant. Only participants with both sample types have a result. For participants with multiple initial or progressive samples, one of each was randomly selected. Only variants that were used in the later study are shown here, although these trends hold when looking at all CBTN variants. GCT: germ cell tumor, ATRT: Atypical Teratoid Rhabdoid Tumor.

The dataset included SVs of five different types: deletions (DEL), duplications (DUP), inversions (INV), insertions (INS), and chromosomal translocations (BND). Overall, BNDs were the most common in the dataset (48%), with DELs second (33%) (**Fig. 1c and Supplementary Fig. 1b**). The distribution of SV types varied across tumor categories. Meningiomas had the highest proportion of DELs at 55% and lowest proportion of BNDs at 33% and metastatic tumors the highest proportion of INVs and DUPs at 12% and 19%, respectively. Samples were extracted from both initial and progressive tumors, with around 75% from initial and 25% from progressive samples (**Fig. 1d and Supplementary Fig. 1c**). As expected, samples from progressive or recurrent tumors had a 1.2 times higher SV burden than ones from initial samples, with 56% of participants having more progressive SVs (**Fig. 1e and Supplementary Fig. 1d**). Interestingly, for some tumor types, including peripheral nerve tumors, the opposite was true, with recurrent tumors having a lower tumor SV burden for 5/6 participants, potentially due to lower heterogeneity in the progressive sample after various selection pressures.

### ML predicts disruptions to 3D genome folding resulting from somatic SVs

Variants from all tumor samples were scored for their effect on surrounding 3D genome contacts using the SuPreMo-Akita pipeline^18^ (**Fig. 2a, Supplementary Table 2, Methods**). In brief, each variant was individually mutated into the reference genome sequence and contact frequency maps for the surrounding 1 Mb region were predicted using Akita. We compared maps corresponding to the reference (REF) and mutated (ALT) genome sequence to measure disruption of genomic contacts using mean squared error (MSE) or 1 - correlation (CORR). A higher MSE or CORR score indicates that a variant is predicted to disrupt genome organization to a greater degree. While we calculate and provide both scores, in some analyses we only focus on one of them for simplicity.

**Figure 2.**
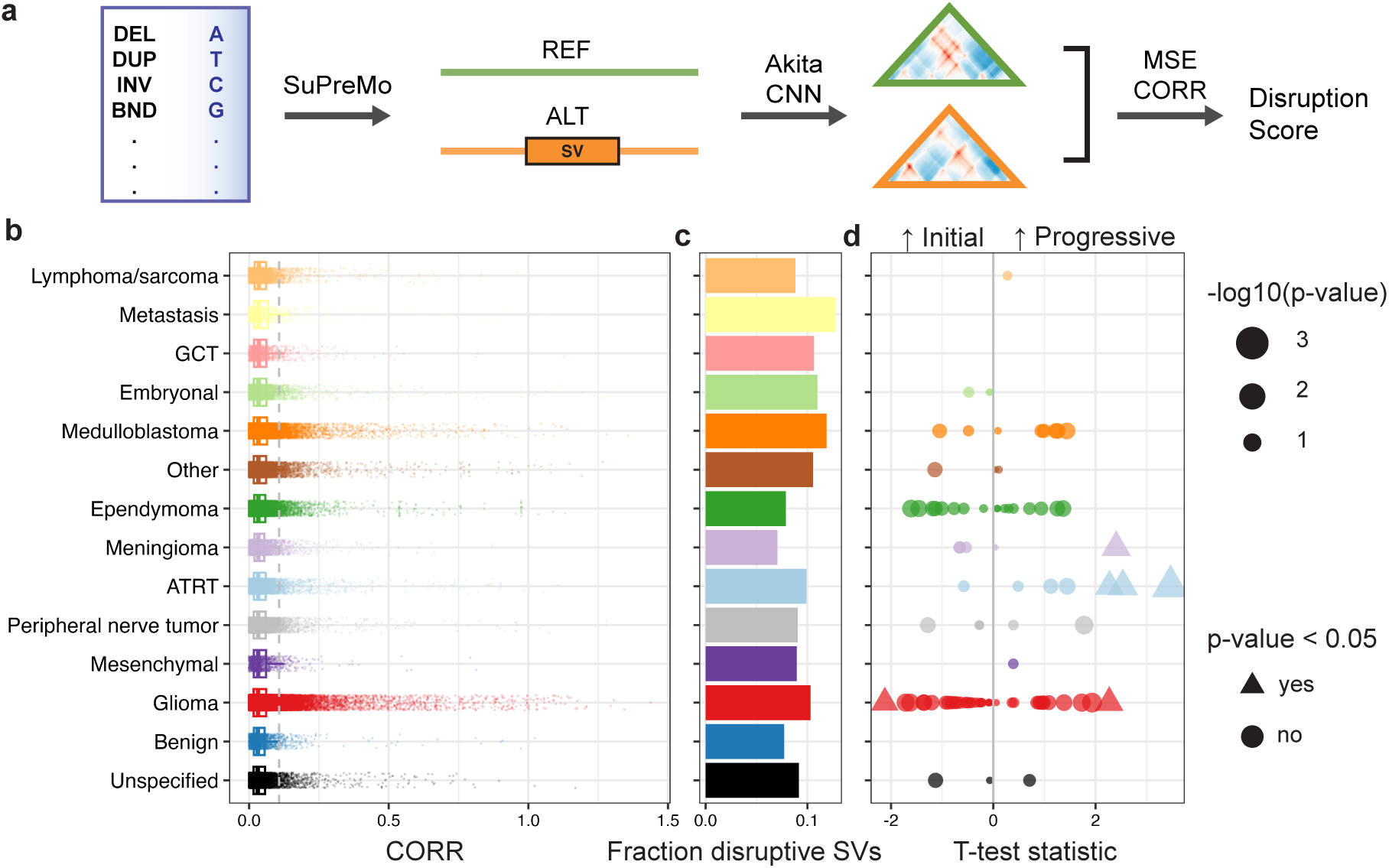
Comparison of 3D genome folding disruption scores. **a.** Schematic for scoring CBTN SVs with SuPreMo-Akita, which takes in variants and outputs disruption scores. It first generates length-matched reference and mutated alternate sequences for each variant. Sequence pairs are inputted into the Akita model which outputs contact frequency maps. The maps are compared using MSE and CORR, which represent the level or disruption that each variant causes to surrounding genomic contacts. **b.** Distribution of disruption scores (CORR) across tumor categories, where each datapoint is a variant. **c.** Proportion of variants that score above the 90th percentile of all CBTN variant scores. **d.** Statistics and *p*-values from t-tests between variant scores from progressive and initial sample pairs (**Methods**). A positive statistic corresponds with progressive SVs being more disruptive than initial SVs. Significant tests (*p*-value < 0.05) are portrayed using triangles while the rest are portrayed as circles.

We compared CORR disruption scores across tumor categories using the median score across all variants in each sample. We found that lymphomas/sarcomas–which are not brain tumors– had the highest median scores across categories (**Fig. 2b**), with rhabdomyosarcomas within this category driving this trend (**Supplementary Fig. 2a**). Germline cell tumors (GCTs) are the highest scoring of the brain tumor categories. Interestingly, benign tumors had the lowest median scores, suggesting that they have higher rates of passenger mutations. Additionally, metastatic tumors had the second highest scores, with the highest proportion of disruptive variants at 13% (**Fig. 2b-c**). This could be because more aggressive tumors have accumulated damaging variants. Neuroblastoma in the embryonal category, and medulloblastoma, both of which have been shown to be associated with genome structure disruptions^12,14,15^, were in the top 5 highest scoring tumor categories (**Fig. 2b**). Most of these trends held up when using MSE disruption scores (**Supplementary Fig. 3a-c**).

We then looked more closely at how tumor types ranked within these categories. For some categories of tumors, like lymphoma/sarcomas, all the included tumor types scored high, in line with the rank of the category overall. For others, like gliomas and embryonal tumors, there was a notable variation that often aligned with how aggressive versus benign the tumor types are. Within the glioma category, brainstem gliomas, gliomatosis cerebri, and oligodendrogliomas scored the highest and gangliogliomas and low grade astrocytomas–both of which are considered to be less aggressive–scored low (**Fig. Supplementary Fig. 2a-b**). Within embryonal tumors, embryonal tumors with multilayered rosettes (ETMR), which are highly aggressive, scored the highest and DNET, which are considered more benign, scored the lowest. Variant effects on genome structure can be visualized in the Akita-predicted contact frequency maps, through the difference between the reference map and the alternate map with the SV and through the disruption track along the map (**Supplementary Fig. 2d**).

While we found that progressive tumor samples had more SVs on average than initial ones, we wanted to evaluate if they also had more damaging SVs. Overall, progressive tumor SVs score significantly higher than initial tumor SVs, with a mean of 0.053 and 0.050, respectively (Welch t-test, p-value = 2.26e-06). For a more direct evaluation, we compared SVs from an initial sample to SVs that arose in a progressive sample from the same individual, which was available for 121/1,843 individuals in the dataset (**Supplementary Table 3**). Overall, progressive samples had higher mean scores than initial samples (**Fig. 2d**), with a mean t-test statistic of 0.1 across all individuals. This difference was significant for 3/7 ATRT participants, 1/2 meningioma participants, and 1/38 glioma participants (**Fig. 2d and Supplementary Fig. 2c**). Higher scores in progressive samples could suggest that they are more aggressive than the initial version of the tumor.

To better understand what makes an SV disruptive to genome structure, we looked at how scores are associated with variant length and location. We found a significant positive association between disruption scores and variant length (**Supplementary Fig. 2e**), as seen for SVs from other disease cohorts^21^. One reason for this might be that larger SVs are more likely to encompass more sequence features that determine genome organization. To account for this, we separated analyses by SV length, when relevant. We also found that there is a relationship between whether the SV overlaps genes or not and how disruptive it is (**Supplementary Fig. 2f**). For non-BND SVs, large coding SVs are more disruptive than large intronic or intergenic SVs. For BNDs–which are all noncoding–intronic SVs are more disruptive than intergenic ones.

### SuPreMo-Akita scores unveil recurrently disrupted genomic regions across tumor types

We next sought to look for patterns in disruption scores along the genome across tumor samples. We evaluated each 1 Mb bin of the genome based on the highest scoring variant for each sample and performed dimensionality reduction. This unveiled groups of samples that clustered separately from the rest due to having disruptive SVs in the same bin. We evaluated these bins and removed low quality predictions (**Methods**). This resulted in five 1 Mb recurrently disrupted regions (RDRs), where clusters of samples had independent somatic SVs that overlapped the region and scored high (**Fig. 3a**). All disruptive SVs in RDRs were DELs and DUPs; some samples had similar SVs overlapping the same RDR while others had a variety of variant lengths and types (**Supplementary Table 4**).

**Figure 3.**
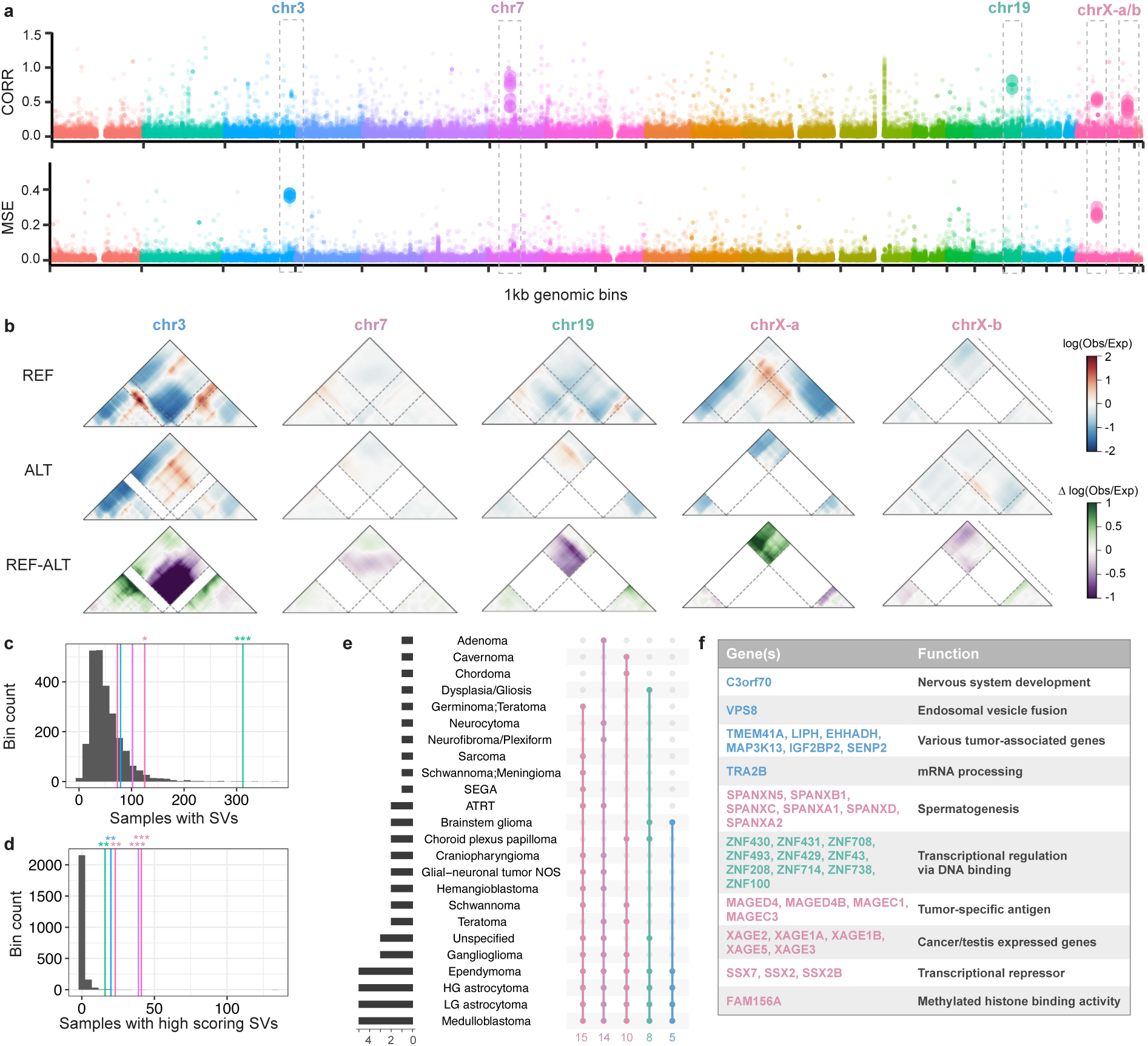
Recurrently disrupted genomic regions. **a.** Disruption scores for the most disruptive SV for each tumor sample across 1 kb genomic bins, ordered by genomic coordinates and colored by chromosome. Sample groups were identified by UMAP on scores for the most disruptive SV (**Methods**). All the clusters had disruptive variants (peaks in the manhattan plot) in the same bin, which are denoted by larger dots. Boxed peaks represent bins that are recurrently disrupted across samples and tumor types using correlation and/or MSE. **b.** Contact frequency maps for example variants from each highlighted bin in a. Reference and alternate maps and the difference between them are shown top to bottom, and dashed lines represent the start and end of the variant. Coordinates for the windows shown, from left to right: chr3:185,492,711-186,410,215, chr7:51,757,685-52,675,189, chr19:20,879,532-21,797,036, chrX:52,172,467-53,089,971, chrX:140,948,793-141,523,916 **c.** Distribution of the number of samples that have an SV at each one of 5 bins. Colored lines represent the number of samples that have an SV in each of the bins highlighted in a. Stars represent significance based on p-value **d.** Same as c but instead looking at the number of samples with an SV that passes the disruption cutoff used in a, for each bin. **e.** Tumor types with variants above disruption cutoff in each of the 5 bins from a, and the overlap across the 5 bins. **f.** Genes in bins from a, grouped by their function.

Changes to surrounding genomic contacts included loss of a TAD boundary in three of the RDRs (chr3, chr7, and chr19) and loss of a loop in another one (chrX-a) (**Fig. 3b**). While only two of the RDRs are mutated in significantly more samples than the rest of the genome (**Fig. 3c**), all 5 are disrupted in significantly more samples than the rest of the genome (**Fig. 3d**), highlighting the benefit of disruption scores in finding these regions. RDRs are disrupted in up to 5-15 different tumor types, with medulloblastoma, low- and high-grade astrocytoma, and ependymoma having disruptive variants in all 5 RDRs (**Fig. 3e**). Interestingly, RDRs include genes involved in nervous system development, transcription, as well as tumor-associated genes, suggesting potential involvement in PBTs (**Fig. 3f**).

To evaluate what makes RDRs stand apart from the rest of the genome, we looked at SV calling confidence and presence of repetitive elements (REs) at SV boundaries. SVs in all RDRs have larger confidence intervals (CIs) than DELs and DUPs in non-RDR bins, suggesting that the exact start and end SV coordinates might be imprecise (**Supplementary Fig. 4a**). Most RDRs have a smaller proportion of SVs with matching REs at their boundaries than the rest of the 1 Mb genomic bins (**Supplementary Fig. 4b**). These mostly include SINE, LINE, simple repeat and LTR elements (**Supplementary Fig. 4c**). The chr19 RDR, on the other hand, has a higher proportion of SVs with matching REs, which might explain why it is recurrently mutated in more samples than other bins (**Fig. 3c**). Overall, these findings lead to the hypothesis that variants in RDRs might play a role in tumorigenesis through disruption of genomic contacts and changes in expression of cancer-causing genes.

### Defining highly disruptive variants

While looking at broad patterns of scores helps us compare tumor types, we only expect that a small subset of variants from this dataset alter genomic contacts that result in transcriptional effects. To help find these, we set a stringent disruption cutoff above which variants result in strong structural changes (**Methods**). This resulted in 363 variants in the dataset that were above this cutoff, which we defined as “highly disruptive”. They were present in only 34/61 of the tumor types. While the majority of high scoring SVs in RDRs described above also fell into the “highly disruptive” classification, we wanted to more broadly evaluate all highly disrupted variants and the genes they might affect. We then defined highly disrupted genes as protein coding genes within 300 kb of highly disruptive SVs that do not overlap their exons. This resulted in 2,012 genes, 249 of which were predicted to be disrupted in more than one tumor type, mostly including medulloblastoma and high-grade astrocytoma (**Supplementary Fig. 5a-b**). These recurrently highly disrupted genes were enriched for myosin complex and axon initial segment (**Supplementary Fig. 5c**).

To focus in on variants that largely have an effect through disruption of genome structure, we isolated variants that passed our strongest disruption cutoff and were noncoding. Out of these 13 variants, 6 resulted in strong changes of contacts of relevant genes (**Supplementary Fig. 6**). For example, a 127,129 base pair (bp) DEL in chromosome 1 removed a TAD boundary and caused gained contacts between the TADs it was insulating, which included *AKT3*, involved in cell proliferation, differentiation, apoptosis, and tumorigenesis and *ZBTB18*, a transcriptional repressor in neuronal development. On chromosome 2, two variants create partial DUPs of TADs that contain *MYCN*–a known tumor associated gene–and *EGLN3*–implicated in renal cell carcinoma^22^–and have the potential to affect their regulation. We hypothesize that these noncoding variants, which do not directly affect genes, might impact the expression of relevant nearby genes by disrupting their distal interactions.

### Disruption is linked to regulatory regions

Chromatin alterations with functional consequences may contribute more to cancer progression, so we focused our analyses on regulatory regions across the genome. To evaluate variant effects on regulatory elements, we identified H3K27ac ChIP-seq peaks and accessible chromatin regions from tumor type-matched cell lines as indicators of potential enhancer activity. We first looked at the association between these regions and variant disruption across tumor types. For pediatric high-grade gliomas (pHGGs), we found that highly disruptive variants were more enriched in H3K27ac peaks and ATAC-seq peaks when compared to the rest of the scored variants, controlled for SV length (**Methods, Fig. 4a**). This trend also held across an additional three tumor types for which we had the corresponding epigenetic data: atypical teratoid rhabdoid tumors (ATRT), diffuse intrinsic pons glioma (DIPG), and medulloblastomas (**Supplementary Fig. 7a-c**). This suggests that there is an interplay between regulatory element contacts and disruptive variants. Motivated by this finding, we extended our SuPreMo-Akita scores to highlight their effect on regulatory elements and to prioritize those variants most likely to disrupt enhancer-promoter contacts.

**Figure 4.**
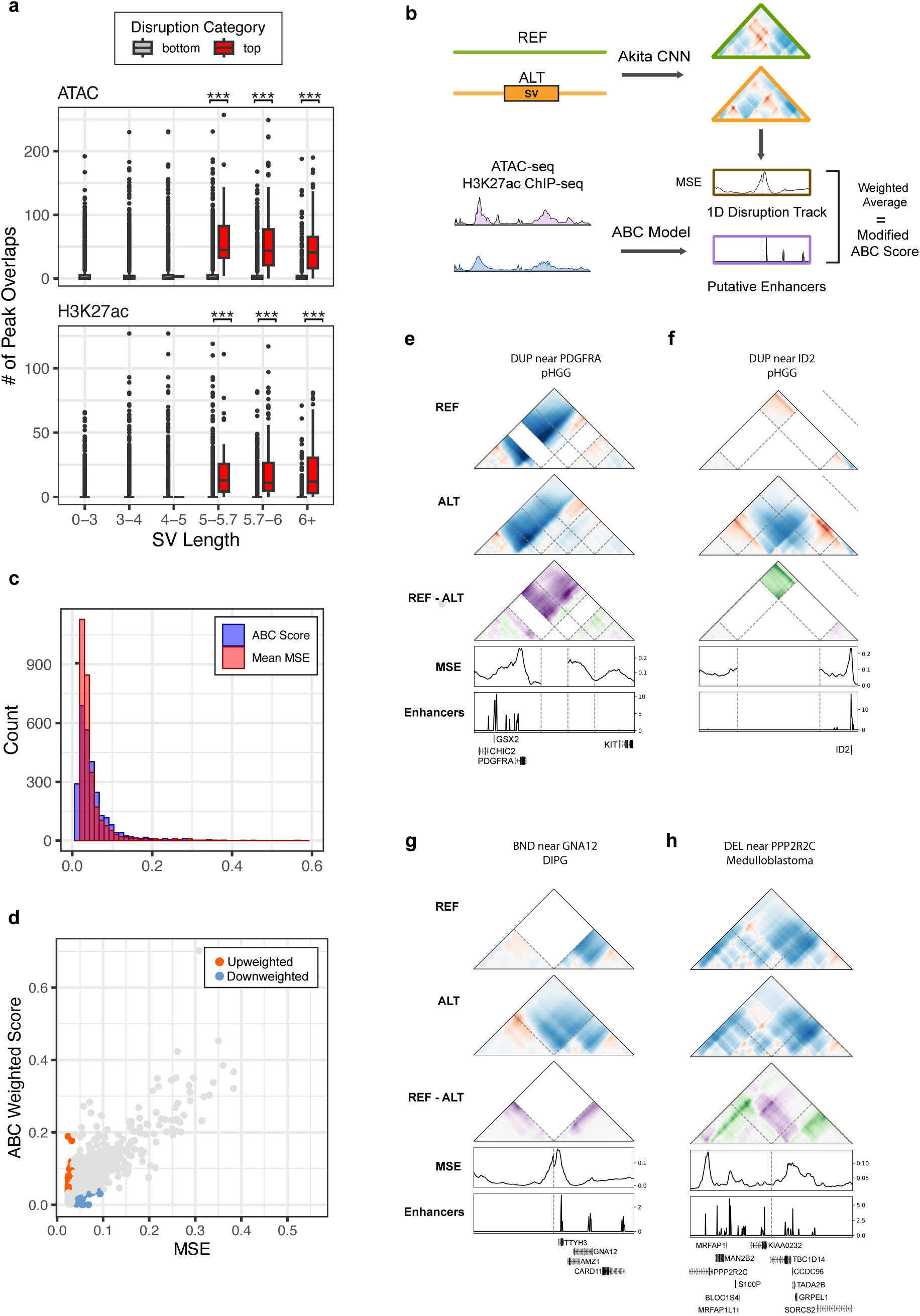
Upscaling disruption at cancer-relevant regions prioritizes previously missed SVs. **a.** Overlap between pHGG SVs with H3K27ac peaks and accessible regions from KNS42, a pediatric glioblastoma cell line. Significant differences in overlap between disruptive and non-disruptive variants at each SV length bin are starred (Mann-Whitney U test, p<0.0001). **b.** Schema to obtain the ABC disruption score, which weights the contribution of predicted 3D genome disruption from an SV based on the enhancer activity in the prediction window. **c.** Histogram of MSE and ABC disruption scores calculated for the top 10% of originally scored pHGG variants. **d.** Scatterplot of MSE versus ABC-weighted scores in pHGG (Pearson *R*=0.8433). Upweighted and downweighted variants are colored in orange and blue, respectively. **e.** Example contact maps of ABC-score prioritized variants near implicated glioma driver genes *PDGFRA* (DUP at chr4:54,365,518 - 54,518,881) and **f.** *ID2* (DUP at chr2:8,498,372 - 8,972,628) in pHGG. **g.** A BND near *GNA12* in DIPG (BND at chr12:115,865,475 - chr7:2,645,728) that was prioritized by ABC disruption scoring. The left and right regions of the horizontal axis for these contact maps depict the 500 Mb region by the breakpoints of the two translocated chromosomes. **h.** ABC-score prioritized DEL near *PPP2RC* in medulloblastoma (DEL at chr4:6,948,839 - 6,949,029).

To upweight variants found at regulatory regions, we took inspiration from the Activity-By-Contact (ABC) Model ^23^, which predicts the strength of enhancer-gene interactions based on information from experimental ChIP-seq, ATAC-seq or DNase-seq, and Hi-C. Using putative enhancers calculated by the model, an “ABC disruption score” for variants from four of the tumor types was determined. In brief, for any loci containing a putative enhancer, the ABC activity score was multiplied by the genome folding disruption score for that region. The final ABC disruption score is a weighted average of the ABC activity score and the 1D disruption score across the 1 Mb region surrounding the variant (**Fig. 4b**, **Methods**). We applied this scoring metric to the top 10% of disruptive variants across pHGGs, DIPGs, ATRTs, and medulloblastomas (**Supplementary Table 5**).

We found that the ABC weighted scores were similar in distribution to and highly correlated with the original disruption scores (**Fig. 4c-d**), suggesting that patterns of genome folding disruption are not skewed towards regulatory regions for most variants. However, we expect that causal variants will drive disruption specifically in functionally important regions, such as enhancers, promoters, and insulating TAD boundaries. In order to identify such variants, we looked at SVs that increased in disruption with the ABC scoring approach compared to the original MSE score (**Supplementary Figure 8a, Methods**). In turn, these upweighted SVs cause focused disruption at enhancer contacts.

The proportion of SV types was consistent across up- and down-weighted SVs, while scores for INVs were largely not changed, likely because of their low scores. There was a trend with SV length: larger SVs were more likely to be downweighted given the smaller remaining region in the prediction map that was evaluated (**Supplementary Fig. 8b-c**). Genes overlapping upweighted variants were found to be involved with the endoplasmic reticulum and immune related gene ontology categories. (**Supplementary Fig. 8d, Supplementary Table 6**). These upweighted variants also affected known cancer genes and genes implicated in cell cycle processes. For pHGG, one of the top three upweighted variants involved *PDGFRA*, a gene encoding a receptor tyrosine kinase that regulates proliferation and survival, and one frequently mutated in gliomas^24^. We saw both coding and noncoding pHGG variants near *PDGFRA* that disrupted chromatin contacts, including a DUP that increased predicted genomic contact for an enhancer at the start of the *PDGFRA* gene (**Fig. 4e**). A different DUP drastically decreased genomic contact at regulatory elements near *ID2*, a transcriptional repressor that regulates cell differentiation and tumor proliferation in pHGG and other cancers (**Fig. 4f**)^25,26^. Furthermore, enhancers in both *PDGFRA* and *ID2* have previously been shown to be targets of recurrent SVs in pHGG^27^, suggesting that the ABC disruption scoring approach identified variants with functionally relevant regulatory roles.

These trends were consistent across other tumor types. Additional upweighted variants included a DEL in a DIPG sample that created a gain of contact upstream of *GNA12*, which has been shown to be upregulated in gliomas (**Fig. 4g**)^28^. *GNA12* activates the RhoA/ROCK signaling pathway, which is involved in cancer cell growth and progression. A DEL in medulloblastoma caused a predicted loss of contact at an enhancer near *PPP2R2C*, a gene involved in synaptic plasticity that has been linked to brain cancers and other mental disorders (**Fig. 4h**) ^29^. Overall, the weighted scoring approach emphasized changes at active enhancer regions, and identified variants that were overlooked with the overall disruption score.

### Upweighted Variants in ATRT Disrupt Candidate Cancer Genes

We placed a greater focus on ATRT, given that it had the highest fraction of high-scoring variants, and highest scoring initial tumor variants. Additionally, ATRT had a slightly larger number of variants with an increase in rank after ABC scoring (**Supplementary Fig. 7d**). ATRT is primarily driven by loss of *SMARCB1*, a subunit of the SWI/SNF chromatin remodeling complex. However, clinical heterogeneity in ATRT is unexplained by the loss of *SMARCB1* alone ^30^, and efforts are ongoing to understand the remaining genetic factors that may contribute to this deadly tumor. To identify possible candidate genes that contribute to ATRT tumorigenesis, we first looked at the most upweighted variants. Several of these variants occurred near genes linked to chromatin organization and *SMARCB1*, such as *CCND1* and *BCL7C* (**Supplementary Fig. 9a-b**). Studies have shown that *CCND1* is a target of 3D genome changes in cancer, and that loss of *SMARCB1* induces *CCND1* deficiency ^31,32^. *CCND1* codes for a component of the CDK complex, which in turn regulates the cell cycle and cell growth. Like *SMARCB1, BCL7C* is a subunit of the SWI/SNF complex, and it has been suggested as a biomarker in gliomas ^33^.

We also highlight a few upweighted variants with corresponding expression changes at nearby genes. All of these variants were BNDs, with high genome folding disruption at putative enhancers of the translocated genes (**Fig. 5a-c**), consistent with enhancer hijacking. These include *NASP*, *MSH6*, and *FOXJ2*. When comparing the samples containing these variants with other ATRT samples containing no variants near these genes, the variant-containing samples had gene expression values more extreme than other samples. The samples with variants near *NASP*, *MSH6*, and *FOXJ2* had expression lower than 31/32 samples, higher than 31/32 samples, and higher than 27/28 samples, respectively. Each of these genes have been implicated in other cancers such as glioblastomas, melanomas, ovarian, and colorectal cancers, and they help regulate processes including histone transport, neural development, DNA damage repair, and cell growth^34,35,36^. We observed similar patterns for several of the other genes near these variants (**Supplementary Fig. 10**), supporting our hypothesis that the SV-driven genome folding alterations may be closely tied to functional effects. Overall, these results demonstrate that ML models can prioritize SVs that disrupt 3D genome folding near neurological and cancer-relevant candidate genes, which help generate new hypotheses regarding their functional roles.

**Figure 5.**
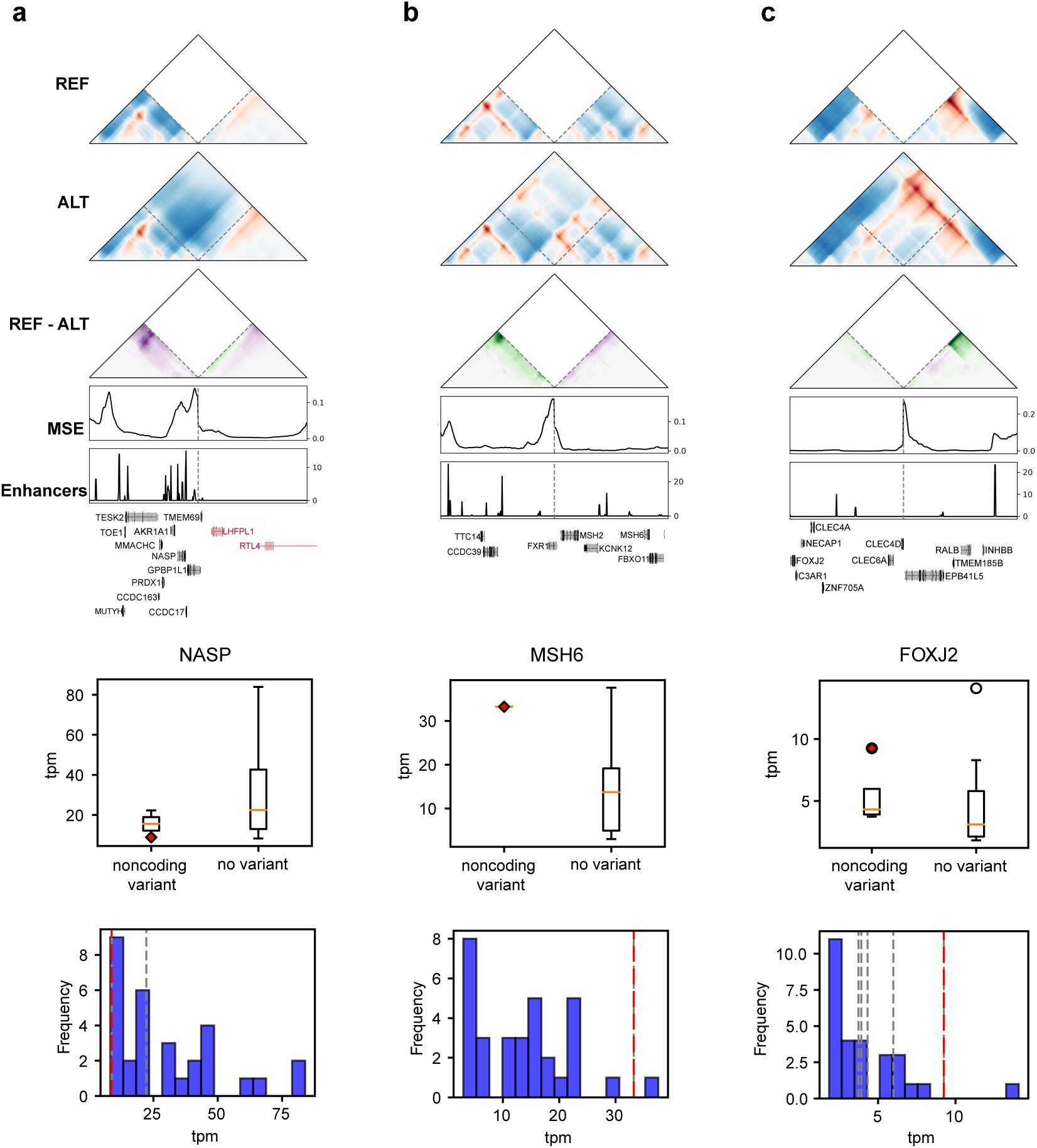
Upscaling disruption at cancer-relevant regions suggests potential new disruptive variants in ATRT. **a-c.** Candidate upweighted BND variants with the ABC disruption score. Disruption is high at putative enhancers near **a**. *NASP* (chr1:45,703,450 - chrX:112,696,292), **b.** *MSH6* (chr3:180,939,147 - chr2: 47,413,958) and **c.** *FOXJ2* (chr12:8,521,326 - chr2:120,042,934). The middle boxplots show gene expression (tpm) for samples containing a noncoding variant versus no variant at all near the gene of interest. The red diamond marks the sample containing the variant shown at the top. The bottom histograms display the distributions of gene tpm values for samples with no variants near the gene. Dotted lines denote the expression values of samples with noncoding variants near the gene. The sample containing the shown variant is marked with a red dashed line.

## Discussion

Here, we present an ML approach to rapidly and systematically screen an unprecedented number of variants to test if they alter 3D genome organization. We applyed this computational approach to characterize somatic SVs across 61 different tumor types from the CBTN cohort. Our data revealed that different tumor types experienced a range of genome folding disruption, with larger SVs and SVs from progressive samples being more disruptive. We found regions that were recurrently disrupted in up to 15 tumor types, which contained genes linked to tumorigenesis and neural system development. Strikingly, several of these 3D disruption hotspots were not mutation hotspots, indicating that SVs in these loci have an elevated propensity to alter genome folding and that our method was essential for identifying their convergent effects. Additionally, we demonstrated that a small number of highly disruptive SVs near cancer genes can be prioritized from a pool of hundreds of thousands of variants, most of which have minimal predicted effects on genome folding. Prioritized SVs were associated with active regulatory regions, leading us to integrate epigenetic data into a new ABC disruption scoring approach to prioritize variants relevant in cancer. Focusing on four tumor types, we identified candidate causal SVs that may contribute to tumorigenesis due to changes in genome structure affecting genes with known and plausible roles in cancer.

Our approach has several limitations, the first of which is the available SV set. Due to their high genomic complexity, SVs often co-occur, and detangling the exact boundaries of each SV can be challenging, especially with short-read sequencing. Hence, there exists a margin of error in the SVs detected and their precise breakpoints. In addition, our screen is dependent on the Akita algorithm and its learned sequence determinants of genome folding. Akita can only make predictions for a 1 Mb window, preventing us from evaluating larger SVs or broader structural changes (e.g. compartment changes). Additionally, Akita was not trained on the specific cell types relevant to pediatric tumors. This is somewhat mitigated by the fact that genome organization is relatively stable across tissues and by our use of epigenetic data from cancer cell lines to upweight chromatin interactions at enhancers in relevant tumor types. Nonetheless, some context-specific chromatin interactions may be missed. Despite these limitations, our predictions serve to generate new, testable hypotheses about hotspot loci and candidate causal variants that can be evaluated in future laboratory studies. To gain further confidence and insights into our results prior to experimentation, they can be combined with predictions from other ML models, such as those predicting chromatin accessibility or transcription factor binding^37^.

WGS is increasingly used in the clinic, and SuPreMo-Akita can contribute to the interpretation of variants beyond obvious gene fusion and inactivation events. The availability of variant data for larger cohorts will necessitate the use of computational strategies, such as ours, to narrow the search space in research projects aimed at elucidating causal mechanisms and identifying new pathways that can be targeted to treat or prevent cancers. Overall, this study shows the utility of ML in identifying functional SVs, and further strengthens previous evidence that SVs contribute to tumorigenesis through changes in genome folding.

## Methods

### Data used

SVs were previously generated by the CBTN^16^ with the Manta SV caller (v1.4.0)^38^. Somatic SVs were called, and only those that passed filtering (‘PASS’) were used. Gene expression data was previously measured and available on the CBTN database in the form of tpm. Variants were previously annotated using AnnotSV^39^. Annotated data was missing for 2/2,386 due to an issue with the vcf file. We grouped tumor types into tumor categories based on their cell types of origin.

### Scoring variants with SuPreMo

SuPreMo-Akita^18^ was used to score all CBTN somatic SVs for the disruption they cause to genomic contacts in the surrounding ∼ 1 Mb. Scores were generated with augmentation, representing the median or mean score among the following sequences: no augmentation, -1 bp shift, +1 bp shift, and reverse complement. MSE and CORR were used to score the differences between the reference and alternate contact frequency maps, with higher scores corresponding to more disruptive variants. Discussion of trends was focused on CORR scores, but most trends were consistent when using MSE (**Supplementary Fig. 3**). Scores only reflect changes to contacts in the map outside of the SV, which is masked from the heatmap. Due to Akita’s sequence length limitation, variants greater than 500 kb in length were removed from the analyses since the remaining portion of the map was deemed insufficient for interpretation. This resulted in 75,073 remaining variants that were scored.

### Recurrently disrupted regions

To find patterns in variant disruption across the genome, we performed dimensionality reduction on disruption scores across all variants and samples. The genome is divided into 1 Mb bins across all chromosomes, and for each bin, only the variant with the highest disruption scores for each sample is considered. This results in two sample-by-bin matrices of maximum disruption scores, one using CORR and one using MSE. We used UMAP to reduce this data to two dimensions which resulted in various small clusters of samples that are separate from the rest. These clusters were samples that all had at least one variant with disruption scores past a cutoff in one bin compared to the rest of the genome. We qualitatively evaluated all variants that overlap each of the selected bins and excluded bins where the variant effects were not strong. We also excluded bins where the predicted chromatin contact map for the reference sequence did not match experimental contact maps due to low quality data, since we are less confident in predictions in these regions. The remaining five bins were considered RDRs and labeled based on their chromosome: chr3, chr7, chr19, chrX-a, and chrX-b. Note that these analyses were performed considering BNDs as individual variants and without excluding initial variants from progressive samples. While most progressive samples have different variants than initial ones, there is some overlap (**Supplementary Table 3**). Additionally, some SVs are repeated across multiple samples from the same individual, although this only occurs in the chr7 RDR.

### Comparing initial and progressive samples

To compare initial samples to progressive samples from the same individual, we took all individuals that had at least one initial and one progressive sample. If more than one sample of each category was present, we randomly chose one of the samples and made sure that we used that chosen sample in all downstream analyses. We then removed all SVs found in the progressive sample that were also present in the initial sample from the same individual, based on the SV coordinates and type. The following analyses were performed using the subsetted progressive sample SVs. To get log fold change in SV count (**Fig. 1e**), we used the total number of SVs in the initial versus progressive sample. To get t-test statistics and p-values (**Fig. 2d**), we performed a Welch’s t-test between scores from the initial and progressive sample.

### Defining disruptive variants and genes

To set a disruption score cutoff for qualifying a variant as disruptive, we used a qualitative visual approach. Cutoffs between the 90th to the 99th percentile, with 1% intervals, of both MSE and correlation were evaluated. At each cutoff, the 10 lowest scoring variants above the cutoff were visualized, five based on MSE and five based on CORR. At the 98th percentile, only 3/10 of the maps had structural differences. Next, cutoffs between the 98.5th and 99.9th percentile, with 0.1% percent intervals, were evaluated. The 98.9th percentile was the lowest cutoff where at least 9/10 maps had easily visualized structural differences, so this was chosen as the disruption cutoff. Using this approach, 363 variants were labeled as disruptive across all samples and variant types. This subset mainly included DELs and did not include any INVs. The most disruptive of the latter were visually inspected to find that only 10/30 resulted in structural changes, similar to 13/30 DELs at the same disruption score percentile. Of note, this cutoff does not signify that variants with lower scores are not structurally different, but rather represents a stringent categorization of variants that are most likely to be disruptive. Genes within 300 kb of either end of each disrupted variant, but not overlapping the variant, were labeled as disrupted genes.

### Epigenetic data processing

Raw ChIP-seq and ATAC-seq data from DIPG007, KNS42, D283 and BT16 cell lines (see Data Availability for accessions) were processed with the ENCODE data processing pipelines (https://github.com/ENCODE-DCC/chip-seq-pipeline2, https://github.com/ENCODE-DCC/atac-seq-pipeline) to produce .bam and peak files.

### ABC-modified scoring

We developed an ABC weighted disruption score, extending SuPreMo using ideas adapted from the ABC Model^23^. In brief, around 150,000 candidate regulatory elements 500 bp in length were identified from H3K27ac ChIP-seq and ATAC-seq data from DIPG007, KNS42, D283 and BT16 cell lines. The Activity score from the ABC model is derived from the geometric mean of read counts of ATAC-seq sites and H3K27ac. Using Akita, a 1D disruption track is generated that represents the MSE between the REF and ALT maps for each of the 448 bins in a given prediction window. Any bin overlapping a 500-bp ABC enhancer is assigned the corresponding ABC Activity score; other bins have a score of 0. For non-BND SVs, the final disruption score is a weighted average of the 1D disruption track and the ABC Activity score track. For BNDs with multiple breakpoints, the final disruption score is the average of all weighted averages of the 1D disruption track and ABC activity track.

### Identifying upweighted variants from ABC-modified scoring

The ABC-modified scoring method was applied to the top 10% of disruptive variants from four tumor types. In order to identify variants that were upweighted, we ranked the variants based on their disruption score, for both the (1) original, unweighted MSE score and (2) ABC-weighted MSE score. We then calculated the difference in rank between the two scores, where a positive value indicates an increase in rank for the ABC-weighted score. The difference in rank was normalized based on the total number of variants in each dataset, and a normalized rank score ≥|40| was used to define upweighted and downweighted variants. This corresponded to <1% (0.54-0.79%) of all variants across these four tumor types.

### Gene ontology analyses

Gene ontology (GO) analyses were performed with clusterProfiler in R. GO was performed using a background set of all genes within 300kb of variants. For recurrently disrupted genes, the 249 genes within 300kb of highly disruptive variants were used. For up- and down-weighted genes, genes in the 300kb region upstream and downstream of upweighted variants from the ABC scoring method were used. For the latter, background genes were near SVs of the same tumor type.

### Data Availability

ChIP-seq for DIPG007 and KNS42 cell lines were obtained from the Gene Expression Omnibus with accession code GSE162976. ChIP-seq for D283 and BT16 were obtained from GSE92585 and GSE174446, respectively. ATAC-seq for D283, DIPG007, and KNS42 cell lines were obtained from GSE92585, GSE229453 and GSE162831, respectively. ATAC-seq for BT16 were obtained from ^40^. SuPremo-Akita scores and other results are available with this paper and its Supplementary Information.

### Code Availability

The code to run SuPreMo-Akita was previously published and can be found at https://github.com/ketringjoni/SuPreMo.

## Acknowledgements

We thank Miguel Brown for his assistance in our collaboration with the CHOP team. We thank Maricruz Rivera for helpful feedback in early phases of the project. We also thank the members of the Pollard, Dahmane, and Resnick labs for helpful discussions.

## Funding

This work was supported by the National Science Foundation Graduate Research Fellowship (S.Z.), NIH Common Fund (grant #R03-DE032505 to A.R., N.D., K.S.P.), the Biswas Family Foundation (K.S.P.), and Gladstone Institutes (K.S.P.).

## Author Contributions

A.R., N.D., and K.S.P. conceived this study. B.Z. and D.M. ran SuPreMo-Akita on CBTN variants. K.G. and S.Z. performed all remaining analyses, prepared figures, and wrote the initial manuscript draft. R.E.Y. suggested analyses and guided results interpretation. K.S.P. and N.D. edited the manuscript. All authors reviewed the final manuscript.

## Competing Interests

The authors declare no competing interests.

## Notes

### Competing Interest Statement

The authors have declared no competing interest.

