## Supplementary Data for "Machine learning-predicted chromatin organization landscape across pediatric tumors"

**CBTN Supplementary Information**

**Supplementary Figures** **2**
Supplementary Fig. 1 3
Supplementary Fig. 2 5
Supplementary Fig. 3 7
Supplementary Fig. 4 8
Supplementary Fig. 5 9
Supplementary Fig. 6 10
Supplementary Fig. 7 11
Supplementary Fig. 8 13
Supplementary Fig. 9. 14
Supplementary Fig. 10. 15
**Supplementary Tables** **16**
Supplementary Table 1. Tumor categories. 16
Supplementary Table 2. Disruption scores. 16
Supplementary Table 3. Progressive samples. 16
Supplementary Table 4. Recurrently disrupted regions. 16
Supplementary Table 5. ABC variant scores. 16
Supplementary Table 6. Upweighted variant GO enrichment. 16

### Supplementary Figures

### Supplementary Fig. 1

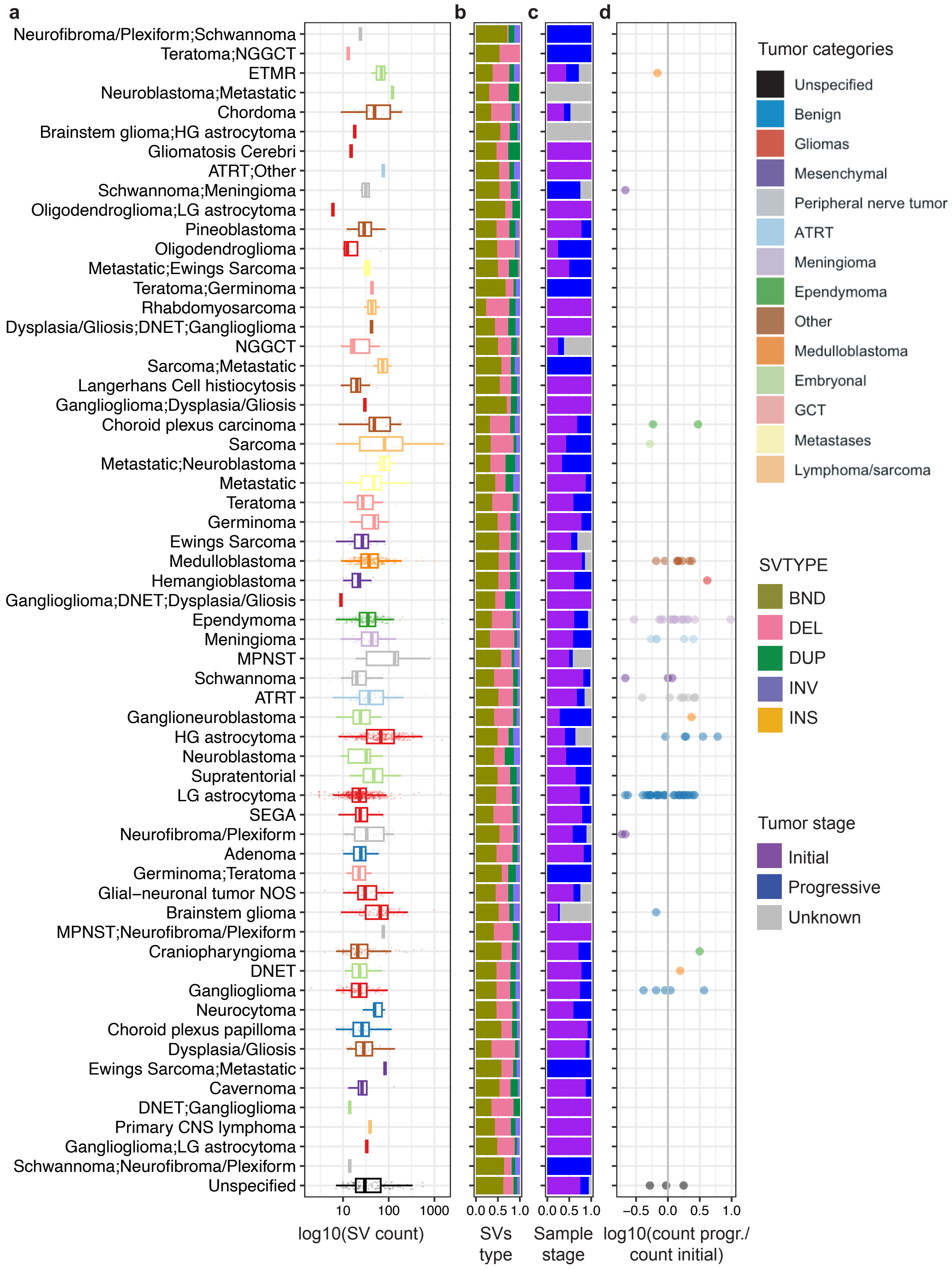

**Supplementary Figure 1. SV profiles across 61 childhood brain tumor types. a.** Distribution of variant lengths across tumor types. Each point is a variant, with multiple samples included in each tumor type. **b.** Fraction of SV types across tumor types out of all SVs from all samples in each type. **c.** Fraction of tumor stages per sample out of all samples in each tumor type. **d.** Log fold change in number of variants in progressive over initial samples from each participant. Only participants with both sample types have a result. For participants with multiple initial or progressive samples, one of each was randomly selected. Only variants that were used in the later study are shown here, although these trends hold when looking at all CBTN variants.

### Supplementary Fig. 2

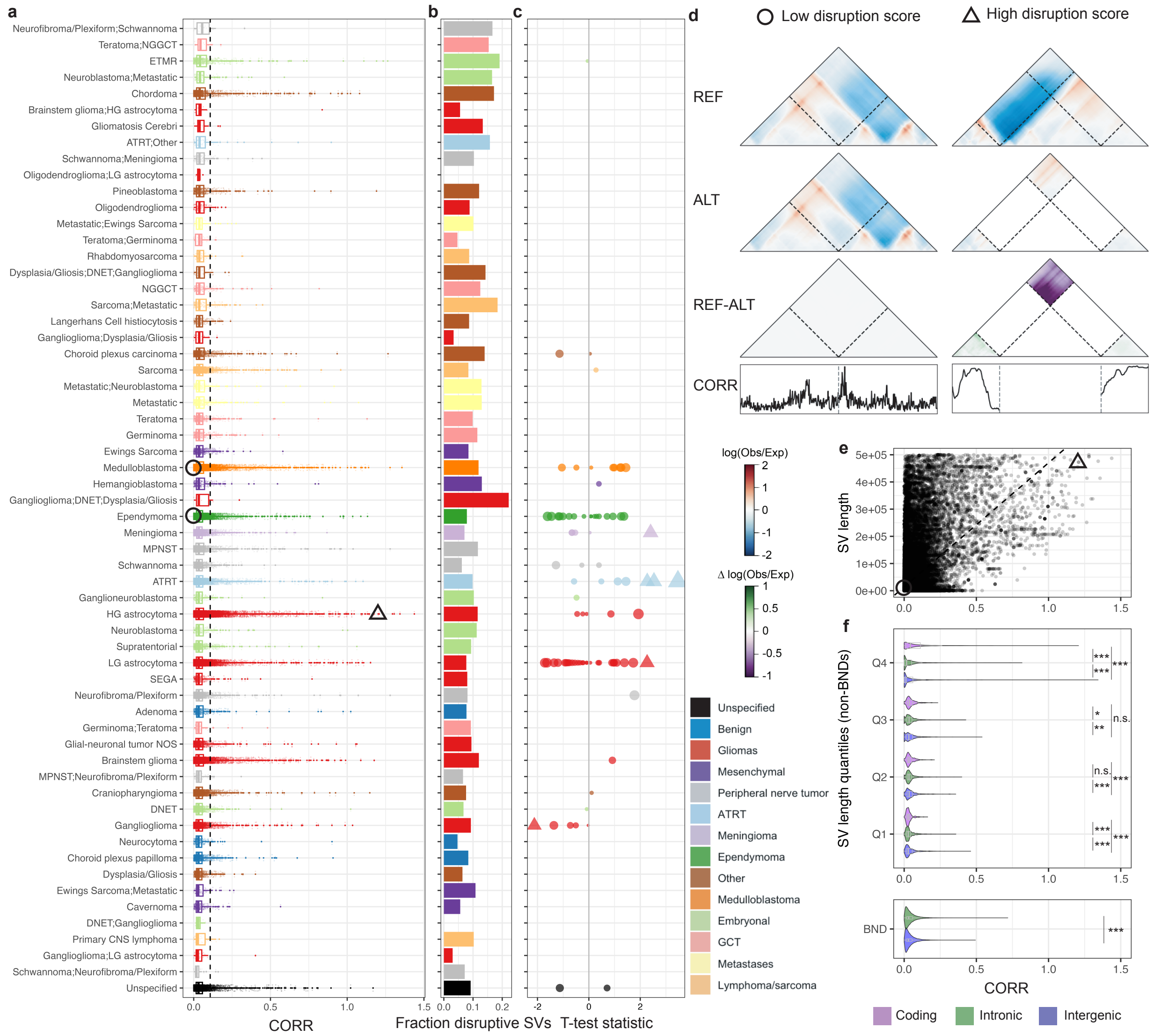


**Supplementary Figure 2. Comparison of 3D genome folding disruption scores across tumor types. a.** Distribution of disruption scores (CORR) across tumor types, where each datapoint is a variant. **b.** Proportion of variants that score above the 90th percentile of all CBTN variant scores. **c.** Statistics and p-values from t-tests between variant scores from progressive and initial sample pairs (Methods). A positive statistic corresponds with progressive SVs being more disruptive than initial SVs. Significant tests (p-value < 0.05) are portrayed using triangles while the rest are portrayed as circles. **d.** Predictions for example low scoring 99 bp INV and high scoring 472,841 DEL. Coordinates for the windows shown: chr2:168,262,816-169,180,320 (left) and chr11:47,875,697-48,793,201 (right). REF: reference sequence; ALT: alternate sequence with SV. **e.** Relationship between SV length and disruption scores. Linear model p-value < 2.2e-16 and adjusted $R^2 = 0.23$. **f.** Distribution of disruption scores across SV locations, separated by SV length quantiles for non-BNDs. SV quantile ranges by quarter are Q1: 0-136, Q2: 137-326, Q3: 327-1,673, Q4: 1,673-499,932 bp. ANOVA test and all Tukey pairwise comparisons have p-values < 2e-16. Non-BND ANOVA p-values are all < 0.01. Tukey HSD pairwise significance shown. BND Mann-Whitney-Wilcoxon Test p-value = 1.019e-11.

**Supplementary Fig. 3**

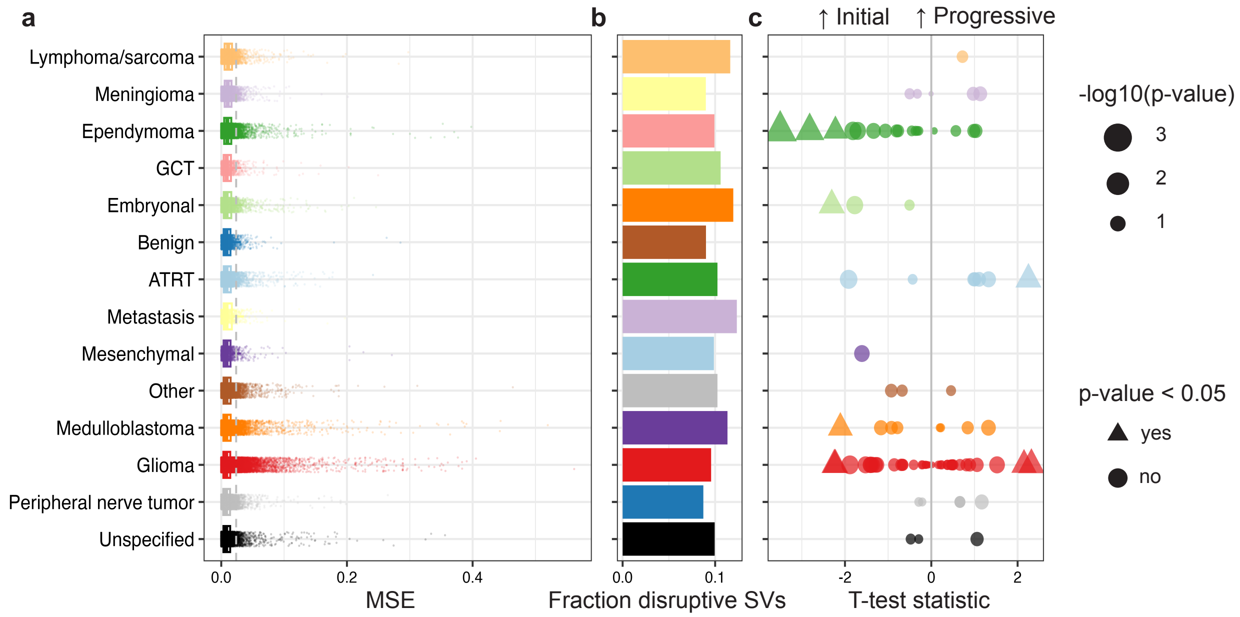


**Supplementary Figure 3. Comparison of 3D genome folding disruption scores using MSE. a.** Distribution of disruption scores (MSE) across tumor categories, where each datapoint is a variant. **b.** Proportion of variants that score above the 90th percentile of all CBTN variant scores. **c.** Statistics and p-values from t-tests between variant scores from progressive and initial sample pairs, generated as in Figure 2 (Methods). A positive statistic corresponds with progressive SVs being more disruptive than initial SVs. Significant tests (p-value < 0.05) are portrayed using triangles while the rest are portrayed as circles.

**Supplementary Fig. 4**

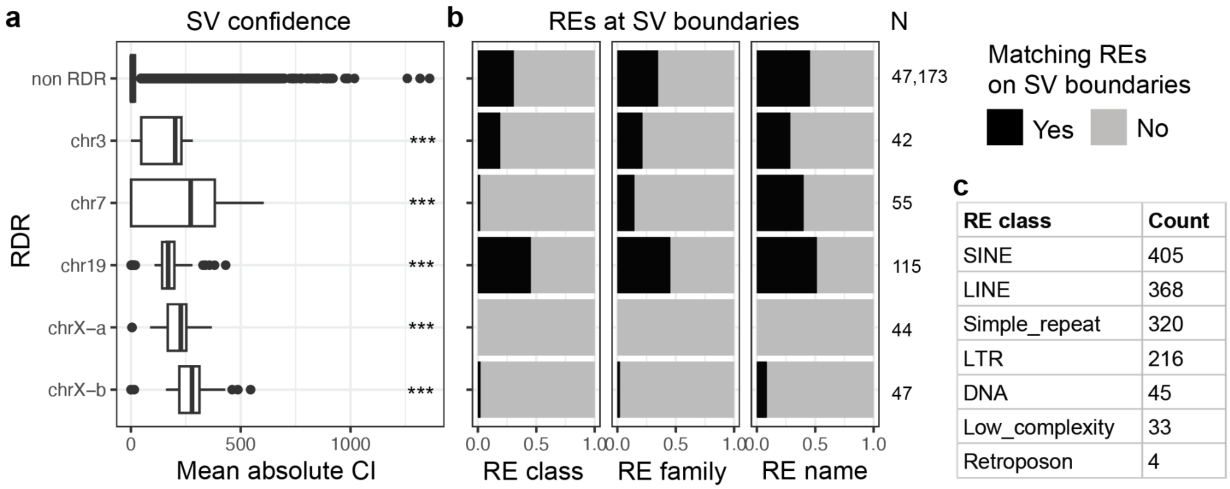


| RE class | Count |
|---|---|
| SINE | 405 |
| LINE | 368 |
| Simple_repeat | 320 |
| LTR | 216 |
| DNA | 45 |
| Low_complexity | 33 |
| Retroposon | 4 |

**Supplementary Figure 4. Confidence and RE presence of SVs in RDRs. a.** Mean absolute confidence interval (CI) at SV start and end positions as generated by Manta, for DEL and DUP in each RDR and in the rest of the genome (non RDR). ANOVA test p-value 0.00095 with Tukey pairwise significance between non RDR and each RDR shown. **b.** Fraction of DELs and DUPs that have matching repetitive elements (REs) within 500 bp of either boundary of the SV, based on the RE class, family, or name. **c**. RE class frequency across RDR SVs. Count represents the number of SVs that have corresponding matching RE at their boundaries. Some SVs have more than one RE class at their boundaries.

**Supplementary Fig. 5**

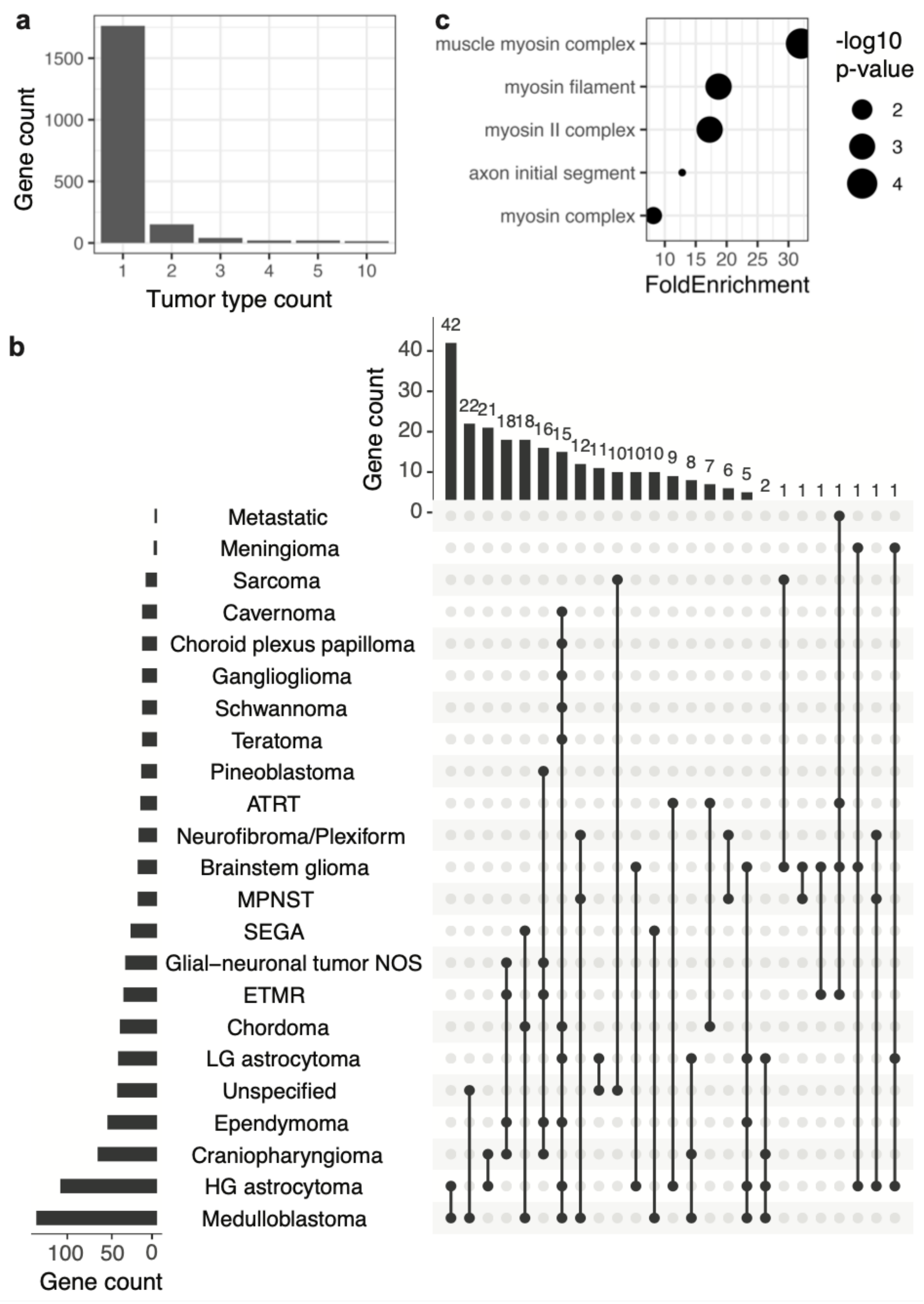


**Supplementary Figure 5. Recurrently and highly disrupted genes. a.** Number of highly disrupted genes and the number of tumor samples in which they are disrupted. **b.** Upset plot of tumor types in which at least 2 genes are highly disrupted. **c.** Enrichment terms of highly disrupted genes affected in at least 2 tumor types.

**Supplementary Fig. 6**

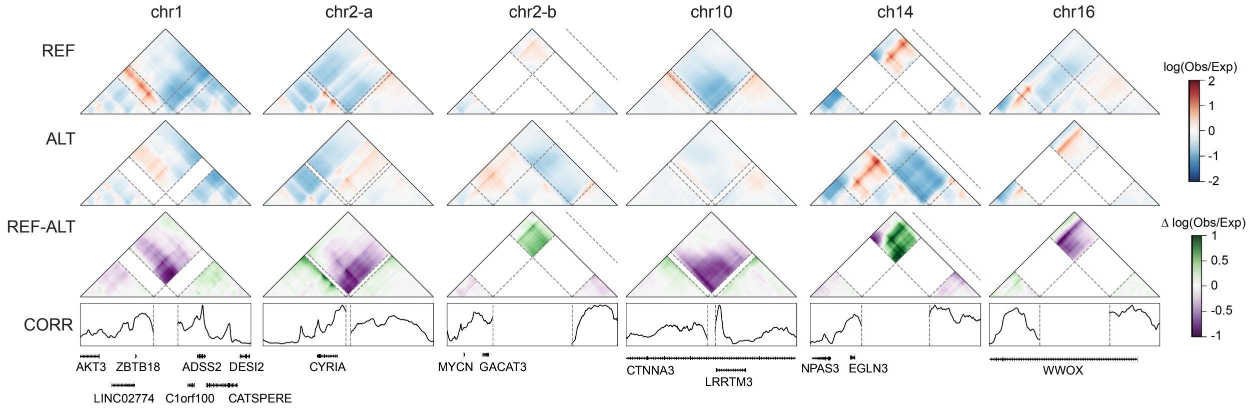


**Supplementary Figure 6. Noncoding variants that disrupt contacts near relevant genes.** Predicted contact frequency maps for the reference, alternate, and the difference shown alongside a track of disruption scores. Coordinates for the windows shown, from left to right: chr1:243,793,865-244,711,369, chr2:16,308,644-17,226,148, chr2:15,863,901-16,356,286, chr10:66,475,920-67,393,424, chr14:33,716,264-34,271,139, chr16:78,427,181-79,344,685.

### Supplementary Fig. 7

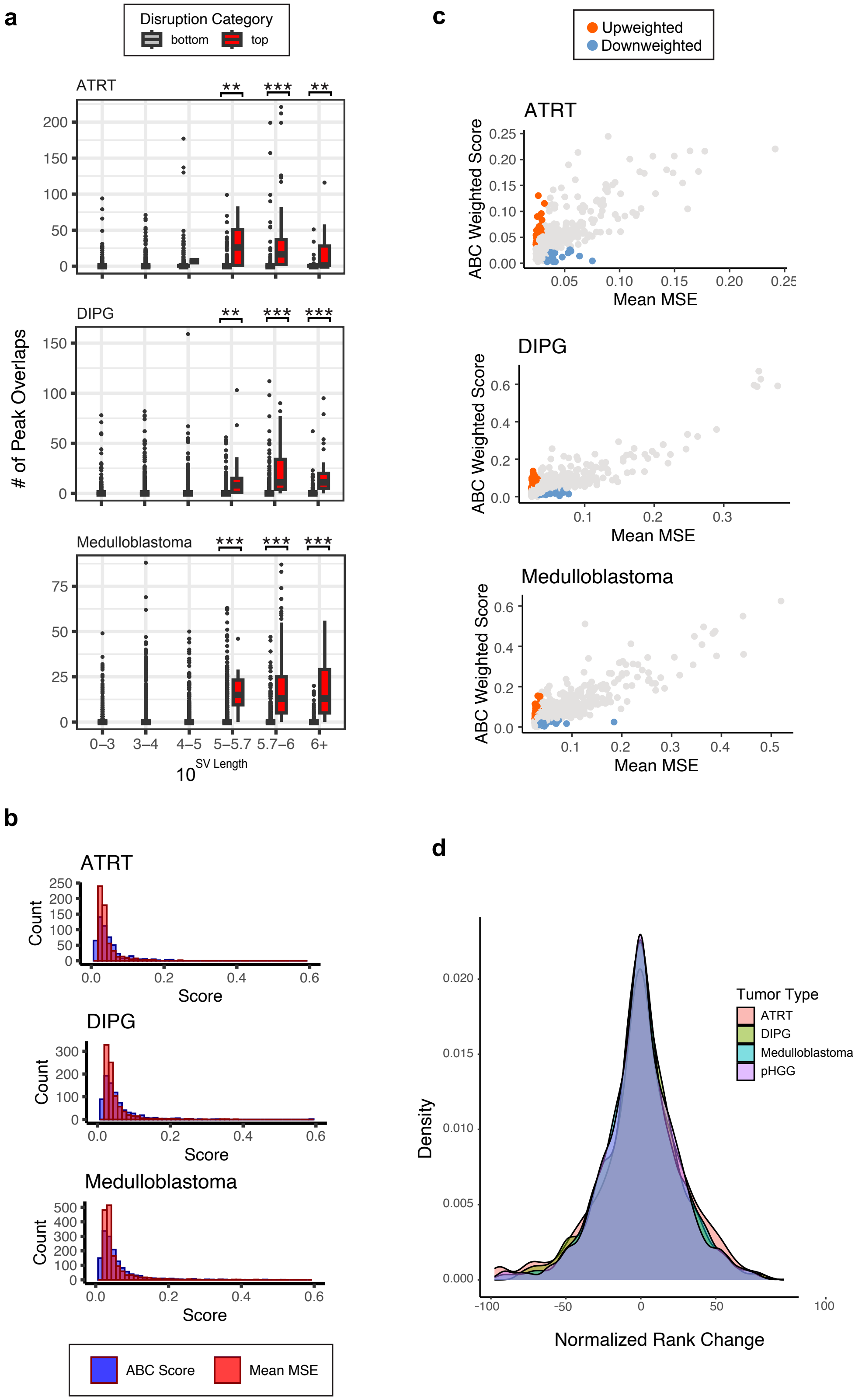

**Supplementary Figure 7. ABC Score Metrics for SVs from ATRT, DIPG, and Medulloblastoma. a.** Overlap between disruptive and non-disruptive SVs from ATRT, DIPG, and medulloblastoma with H3K27ac ChIP-seq peaks and ATAC-seq peaks from tumor-type matched cell lines. **b.** Histogram of MSE and ABC scores calculated for the top 10% of originally scored pHGG variants. **c.** Scatterplot of MSE vs ABC disruption scores for each tumor type. Upweighted and downweighted variants are colored in orange and blue, respectively. **d.** Distribution of normalized change in variant rank across all four tumor types, when going from the MSE to the ABC disruption scores.

**Supplementary Fig. 8**

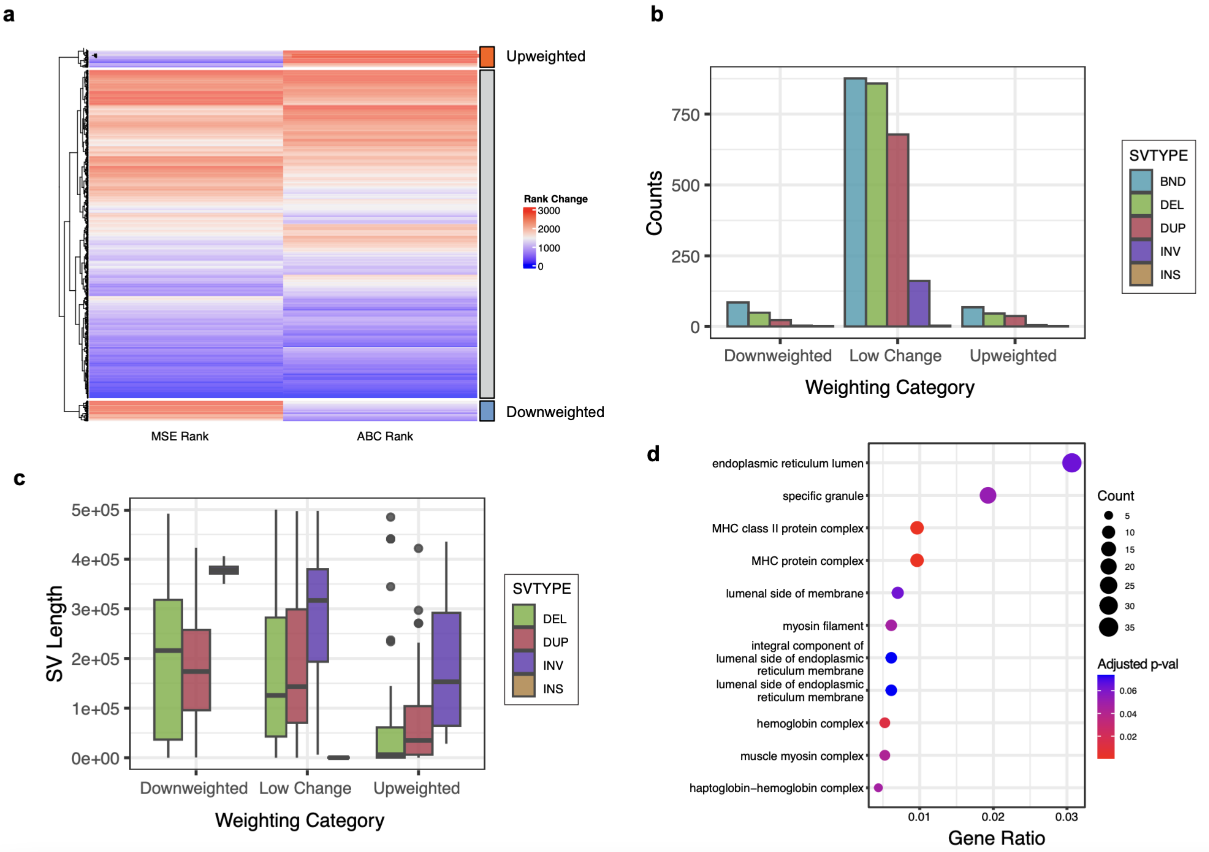


**Supplementary Figure 8. Characteristics of variant disruption in ABC-scored pHGG SVs. a.** Heatmap of SV disruption ranks when scored by MSE vs. the ABC disruption score. SVs are grouped by their rank change. **b.** Number of SVs that were upweighted, downweighted or had a low change in rank based on the ABC disruption score, when compared to MSE. SVs are grouped by SV type. **c.** Distribution of SV lengths for upweighted, downweighted and “no change” SVs. SVs were grouped by SV type, with BNDs removed. **d.** Top scoring GO enrichment terms for genes within 300kb of ABC-upweighted variants.

Supplementary Fig. 9

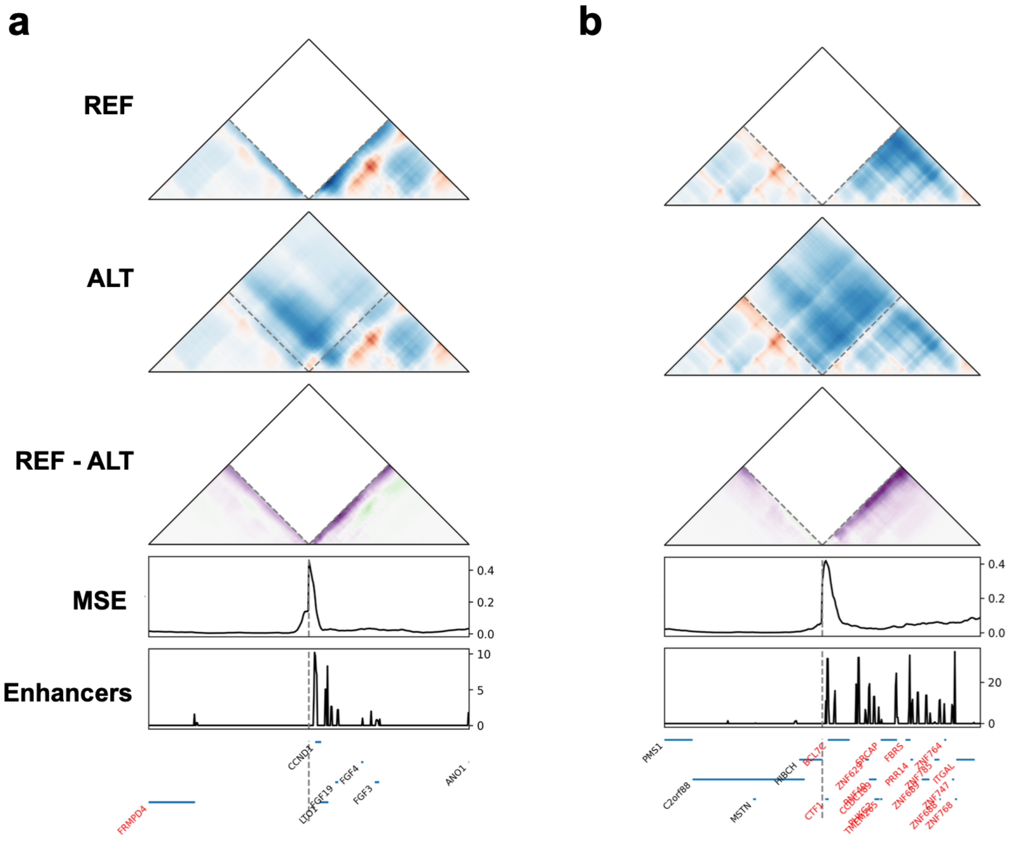


**Supplementary Figure 9. ABC-upweighted variants in ATRT are near genes associated with chromatin remodeling.** Upweighted variants in ATRT with the ABC disruption score. Disruption is high at putative enhancers near **a.** *CCND1* (BND at chrX:11743616 - chr11:69622787) **b.** *BCL7C* (BND at chr2:190323979 - chr16: 30911237). The left and right regions of the horizontal axis for these contact maps depict the 500Mbp region by the breakpoints of the two translocated chromosomes.

**Supplementary Fig. 10.**

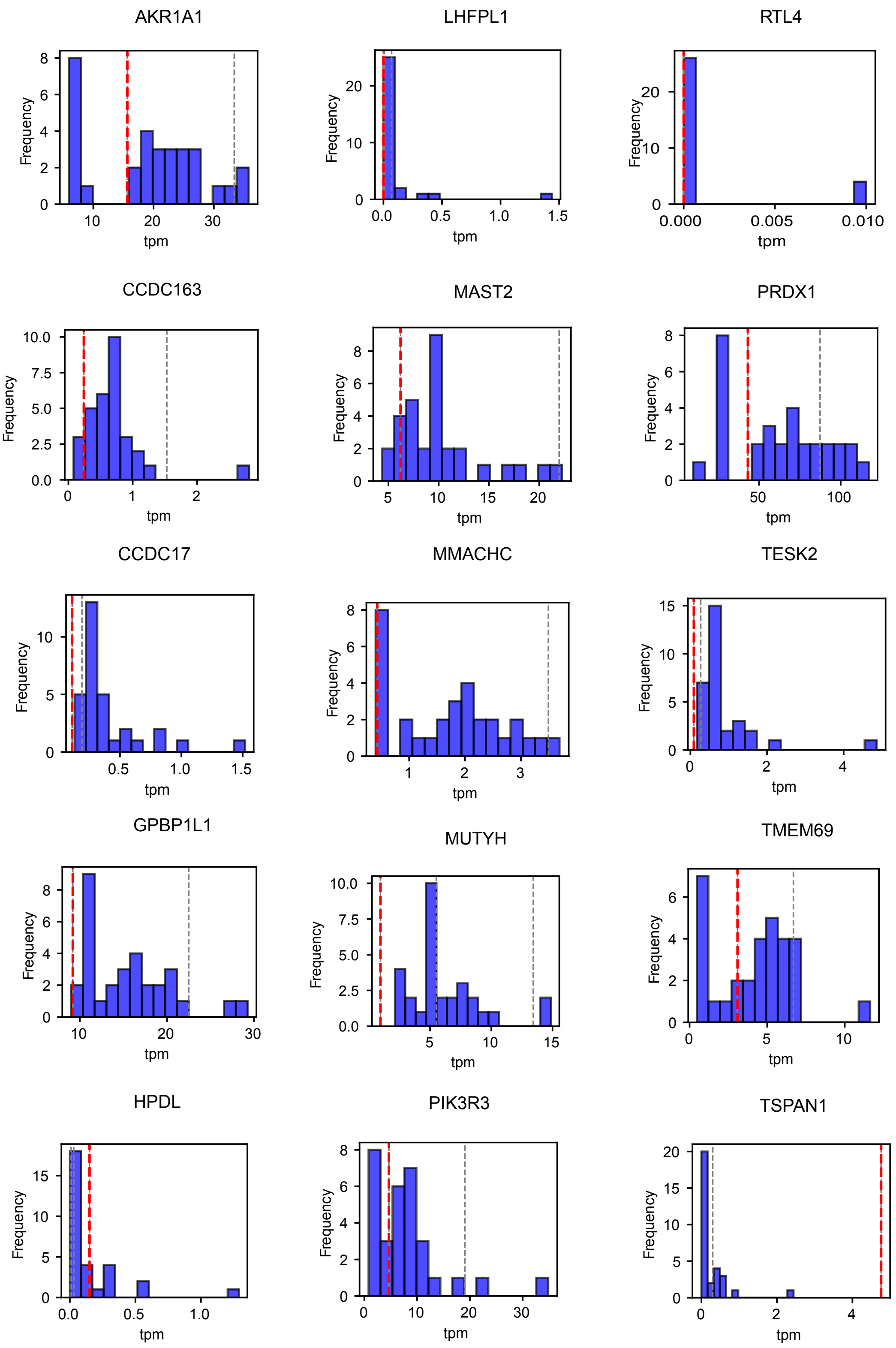

**Supplementary Figure 10. Expression is broadly different for genes near ATRT upweighted variants.** Histograms of gene expression for genes in the prediction window of the BND in **Figure 5a**. Each histogram shows the distributions of expression values (tpm) for samples with no variants near the gene of interest. The dotted lines denote the expression values of samples with noncoding variants near the gene. The sample containing the variant from **Figure 5a** is marked with a red dashed line. Across all samples, the variant-containing sample in red had a tpm value that was in the top or bottom 10% samples for 8/15 of the genes shown.

### Supplementary Tables

**Supplementary Table 1. Tumor categories.**

**Supplementary Table 2. Disruption scores.**

**Supplementary Table 3. Progressive samples.**

**Supplementary Table 4. Recurrently disrupted regions.**

**Supplementary Table 5. ABC variant scores.**

**Supplementary Table 6. Upweighted variant GO enrichment.**
